# Effects of agrochemicals on disease severity of *Acanthostomum burminis* infections (Digenea: Trematoda) in the Asian common toad, *Duttaphrynus melanostictus*

**DOI:** 10.1101/165795

**Authors:** Uthpala A. Jayawardena, Jason R. Rohr, Priyanie H. Amerasinghe, Ayanthi N. Navaratne, Rupika S. Rajakaruna

## Abstract

**Background:** Agrochemicals are widely used in many parts of the world posing direct and indirect threats to organisms. Xenobiotic-related disease susceptibility is a common phenomenon and a proposed cause of amphibian declines and malformations. For example, parasitic infections combined with pesticides generally pose greater risk to both tadpoles and adult frogs than either factor alone. Here, we report on experimental effects of lone and combined exposures to cercariae of the digenetic trematode *Acanthostomum burminis* and ecologically relevant concentrations of (0.5 ppm) four pesticides (insecticides: chlorpyrifos, dimethoate; herbicides: glyphosate, propanil) on the tadpoles and metamorphs of the Asian common toad, *Duttaphrynus melanostictus*.

**Results:** All 48 cercaraie successfully penetrated each host suggesting that the pesticides had no short-term detrimental effect on cercarial penetration abilities. When the two treatments were provided separately, both cercariae and pesticides significantly decreased the survival of tadpoles and metamorphs and increased developmental malformations, such as scoliosis, kyphosis, and skin ulcers. Exposure to cercariae and the two insecticides additively reduced host survival. In contrast, mortality associated with the combination of cercariae and herbicides was less than additive. The effect of cercariae on malformation incidence depended on the pesticide treatment; dimethoate, glyphosate, and propanil reduced the number of cercarial-induced malformations relative to both the control and chlorpyrifos treatments.

**Conclusions:** These results show that ecologically relevant concentrations of the tested agrochemicals had minimal effects on trematode infections, in contrast to others studies which showed that these same treatments increased the adverse effects of these infections on tadpoles and metamorphs of the Asian common toad. These findings reinforce the importance of elucidating the complex interactions among xenobiotics and pathogens on sentinel organisms that may be indicators of risk to other biota.

## Background

Amphibian populations in many parts of the world are experiencing declines and malformations owing to multiple causes, such as xenobiotics, diseases, radiation, habitat destruction, and climate change. [1,2]. Among these causes, considerable attention has been paid to the effects of chemical contaminants on disease risk [3–7]. Amphibians prefer to live in littoral zones of wetland or aquatic ecosystems where there is a high potential exposure to agrochemicals [8] as the pesticides can travel over large expanses as high as 1,000 km affecting it life cycle stages [9, 10].

Many studies conducted on effects of xenobiotic on amphibians have focused on direct mortality and developmental defects that might contribute to population declines [12–15]. For instance, the direct mortality of late stage larvae of green frogs (*Rana clamitans*) and spring peepers (*Pseudacris crucifer*) was studied by exposing them to 3 ppb of the insecticide carbaryl [16]. Relyea *et al*. [12] reported that exposure to 380 ppb of the herbicide glyphosate (Roundup) resulted in a 40% decline in the survivorship of American toad (*Bufo americanus*), leopard frog (*Rana pipiens*), and gray tree frog (*Hyla versicolor*) tadpoles. Other than effects on survival, growth reductions due to pesticide exposure can potentially reduce population growth rates of amphibians [12]. Furthermore, pesticides may delay or accelerate amphibian metamorphosis [17–19]; delays could cause mass mortality events if the waterbody dries up before metamorphosis and accelerated metamorphosis can compromise the immune capacity of metamorphs [20]. In addition to this indirect effect on immunity, pesticides can also be directly immunotoxic increasing susceptibility to infectious diseases [21].

Infectious diseases are particularly important because they are well-documented, widespread causative agents of amphibian population declines [22–24]. Among the amphibian infectious diseases, those caused by trematode infections have received much interest [23, 25, 26]. Deformed amphibians and associated mass mortality events became a major environmental issue during the late 1990’s [4] and later on, trematode infections were identified as the major cause of many of these deformities [27–30]. By deforming their hosts, the trematodes are believed to enhance the chances that the intermediate host is depredated by a vertebrate definitive host, thus facilitating their life cycle completion [the handicapped frog hypothesis; 4, 31].

Agrochemicals consistently seem to affect interactions between amphibian hosts and trematode parasites [4]. For example, *Echinostoma trivolvis* infection of cricket frogs has increased in areas with detectable levels of herbicides in Midwestern United States [33]. Similarly, *E. trivolvis* infection in *Rana clamitans* has increased in areas closer to nutrient and where other chemical inputs were high [26]. To corroborate these findings, Rohr and colleagues [34] demonstrated that the trematode infections were higher in amphibians exposed to atrazine, glyphosate, carbaryl, and malathion. Furthermore, elevated levels of nitrogen and phosphorous associated with fertilizer use increased amphibian trematode infections [34, 36].

In this study, we examined the effects of *Acanthostomum burminis* infections in the tadpoles and metamorphs of the Asian common toad, *Duttaphrynus melanostictus* in the presence of four pesticides: two herbicides (glyphosate and propanil), and two insecticides (chlorpyrifos and dimethoate). Individual effects of these pesticides on *A. burminis* infections in the same developmental stages of the hourglass tree frog, *Polypedates cruciger*, and *D. melanostictus* were previously reported [36–40]. In these species, *A. burminis* induced mainly axial and some limb malformations, increased mortality and time to metamorphosis, and decreased size at metamorphosis [36, 37, 40], whereas the four pesticides increased malformations, mortality, and time to metamorphosis [38, 39]. Many laboratory studies suggest that in the presence of pesticides, trematode-induced effects are enhanced [41–44]. Recently, exposure to the combination of cercariae of *A. burminis* and pesticides revealed that the two factors pose greater risks to frogs than either factor alone [45]. Despite earlier findings that cercariae are often sensitive to chemical contaminants [46–48], all the cercariae entered the tadpole in both the control and pesticide treatments, indicating that there was no pesticide-induced mortality of the cercariae before they could infect [45]. Whereas previous work on *D. melanostictus* explored the effects of pesticides and *A. burminis* in isolation only, here we build upon work that suggests that pesticide exposure can enhance trematode infection by crossing the presence and absence of pesticides with the presence and absence of *A. burminis* to test whether pesticides increase or decrease risk from this infection in *D. melanostictus*. Consequently, this work will help move the field towards a more general conclusion regarding the risk that the combined effect of the pesticides and cercariae pose to amphibians.

## METHODS

### Study animals

The Asian common toad, *Duttaphrynus melanostictus* is a least concerned species, distributed all over Sri Lanka, especially in human-altered habitats. The adults lay egg strands in slow-flowing streams or in water pools. Four newly spawned egg masses of *D. melanostictus* were collected from ponds in the Peradeniya University Park (7°15′15″N 80°35′48″E / 7.25417°N 80.59667°E) and were brought to the research facility in the Department of Zoology, University of Peradeniya, Sri Lanka. The egg masses were placed in a glass aquarium containing dechlorinated tap water. Tadpoles were fed ground fish flakes twice a day (~10% body mass). The debris and faeces that collected at the bottom of the aquaria were siphoned out and water level was replenished daily. Water temperature was maintained around 27º −30º C and pH was maintained around 6.5-7.0.

Adults of *Acanthostomum burminis* reproduce sexually in the common freshwater snake [40] and release eggs in the excrement of these hosts. Miracidiae, a free-living larval stage, hatch when the eggs encounter water and search for the first intermediate host, a snail. Once in the snail host, they reproduce asexually and a second free-living larval stage, cercaria, is released. Cercariae search for their second intermediate host, which is an amphibian. The cercariae encyst subcutaneously as metacercariae in amphibians. When an infected amphibian is consumed by a water snake, the life cycle is completed.

Pleurolophocercous cercariae of *A. burminis* released from the freshwater snail species *Thiara scabra* (Family: Thiaridae) were used for the trematode exposures in this experiment. *Thiara scabra* is a common freshwater snail, found associated with muddy/sandy bottom closer to riverine vegetation [49]. *Thiara scabra* were collected from the university stream and were kept in plastic vials containing 10–15 mL of dechlorinated tap water, under sunlight to induce cercarial shedding. The snails that were shedding cercariae were kept individually in separate vials to obtain a continuous supply of cercariae for the exposures. One infected snail was used for all the tadpole exposures per clutch. Thus, four source snails were used to expose the tadpoles from the four clutches of toads. This is advantageous because the blocking factor removes variation from the error term that is due to both the source of the tadpoles (clutch) and the source of the cercariae (snail), increasing statistical power to detect an effect of treatments.

### Test chemicals

The tadpoles and cercariae were exposed to commercial formulations of four widely used agrochemicals; two organophosphorous insecticides (chlorpyrifos and dimethoate) and two herbicides (glyphosate and propanil). The concentration of the active ingredient (a.i.) tested and any known surfactants in the commercial formulation were given in Table 1. The test concentrations (0.5 ppm) for each pesticide were selected based on available literature [50, 51] and information from Pesticide Registrar Office in Peradeniya on field concentrations of these chemicals.

**Table 1.** Active ingredient, surfactant and commercial name of the four pesticides used

| Active ingredient and strength | Surfactant/solvent | Trade name |
| --- | --- | --- |
| Chlorpyrifos[O,O-DiethylO-(3,5,6-trichloro-2-pyridyl) phosphorothioate] 400g/L | Xylene | Lorsban 40 EC<sup>®</sup> or Pattas<sup>®</sup> |
| Dimethoate (O,O-Dimethyl phosphorodithioate) 400 g/L | Water | Dimethoate 40EC<sup>®</sup> |
| Glyphosate [N-(Phosphonomethyl) glycine] 360 g/L | POEA | Round Up<sup>®</sup> or Glyphosate<sup>®</sup> |
| Propanil [N-(3,4-dichlorophenyl) propanamide] 360 g/L | Cyclohexanone & petroleum solvents | 3, 4-DPA<sup>®</sup> |

### Exposure of tadpoles to ceracraie and pesticides

Each tadpole (5 days post-hatch, Gosner stages 25–26 [52]) was placed in a separate specimen cup containing 15–20 mL of test solution (dechlorinated tap water/0.5 ppm-chlorpyrifos/dimethoate/glyphosate/propanil). Tadpoles assigned to receive trematodes (Table 2) were exposed to12 cercariae per day for four consecutive days. Cercarial penetration was observed under a dissecting microscope and the containers were examined every half hour to ensure that no free swimming cercariae remained. A total of 800 tadpoles were tested requiring 20 randomly selected tadpoles from each clutch for each treatment (20 tadpoles per clutch × 5 pesticide treatments × 2 trematode treatments ×4 clutches = 800 tadpoles). After exposure to the cercariae, 20 tadpoles of each treatment regime were assigned to one of 10glass aquaria (15 x 15 x 25 cm) containing 2 L of one of the five test solutions (dechlorinated tap water or 0.5 ppm of chlorpyrifos, dimethoate, glyphosate, or propanil). The tadpoles were raised in the same test medium until metamorphosis. The test solution was renewed once a week and temperature and pH were maintained between 26 and 30° C and 6.5 and 7.0, respectively under a natural photoperiod of approximately 12:12 h.

**Table 2.** Experimental design used to test the individual and combined exposure of *Acanthostomum* cercariae and pesticides on *D. melanostictus* tadpoles

| Parasite | Exposure medium |  |  |  |  |
| --- | --- | --- | --- | --- | --- |
|  | Dechlorinated tap water (Control) | Chlorpyrifos | Dimethoate | Glyphosate | Propanil |
| No cercariae | - | - | - | - | - |
| Cercariae<br>(12×4=48) | + | + | + | + | + |

Note: Concentration of 0.5 ppm was used for all four chemicals. 20 tadpoles were tested in each treatment

### Data collection and analyses

Tadpole mortality, forelimb emergence (stage 42, [52]), and metamorphosis were assessed daily. When dead tadpoles were noticed, they were removed and preserved in 70% alcohol. Snout vent length (SVL) to nearest 0.01 cm and body mass to nearest 0.001 g were recorded at metamorphosis. Malformations were reported at 10 and 30 days post hatching for larvae and at metamorphosis. Malformations were identified and categorized according to Meteyer [53], and severely malformed metamorphs were euthanized with MS-222 and preserved. All procedures described herein were approved by the Animal Ethical Review Committee (AERC/06/12) at the Postgraduate Institute of Science, University of Peradeniya.

Data were analyzed using Statistica (version 6) software (Statsoft, Tulsa, OK). We used binomial regression to test whether temporal block and the main and interactive effects of pesticide and cercarial treatments affected the proportion of frogs that survived in each tank. The binomial error distribution was further used to assess how the treatments affected malformation frequency of 10 days post-hatching. We used a general linear model to test whether the effect of temporal block and the main/interactive effects of pesticide and cercarial treatments affected the SVL, mass at metamorphosis, TE50 [day that 50% of the animals in a tank had forelimb emergence-stage 42], and days to metamorphosis. Because all 20 tadpoles were reared in a single tank in each temporal block, we used the mean of each tank as the replicate and thus each treatment had four replicates total for these analyses. If any main effect or interaction were significant, a Fisher’s least significant difference (LSD) Posthoc test was conducted to evaluate which treatments differed from one another. If temporal block was not significant, it was dropped from the statistical model.

## RESULTS

All the cercariae penetrated each tadpole because none remained in the vials at the end of exposure. Hence, attempted infections appear to be the same across pesticide treatments.

### Survival of the tadpoles

Both the effect of pesticide treatment (Main effect: χ^2^_4_= 54.53, *p*= 0.0001) and cercariae (Main effect: χ^2^_1_= 46.31, *p*= 0.0001; Fig 1a) increased tadpole mortality (Fig. 1A). In the absence of cercariae, all four pesticides significantly increased mortality (Fig. 1A). Tadpoles exposed to chlorpyrifos, dimethoate, glyphosate, and propanil had 6.50, 7.75, 7.75, and 4.25 times the mortality as those exposed to the pesticide control (Fig. 1A). In the absence of pesticides, tadpoles exposed to cercariae had 6.75 times the mortality as those not exposed to cercariae (Fig. 1A). There also was an interaction between pesticide and cercarial treatments on the probability of death (χ^2^_1_= 11.02, *p*= 0.026, Fig. 1A). This interaction was caused mostly by the combination of herbicides and cercariae having a less than additive effect on mortality (Fig. 1A).

**Figure 1.**
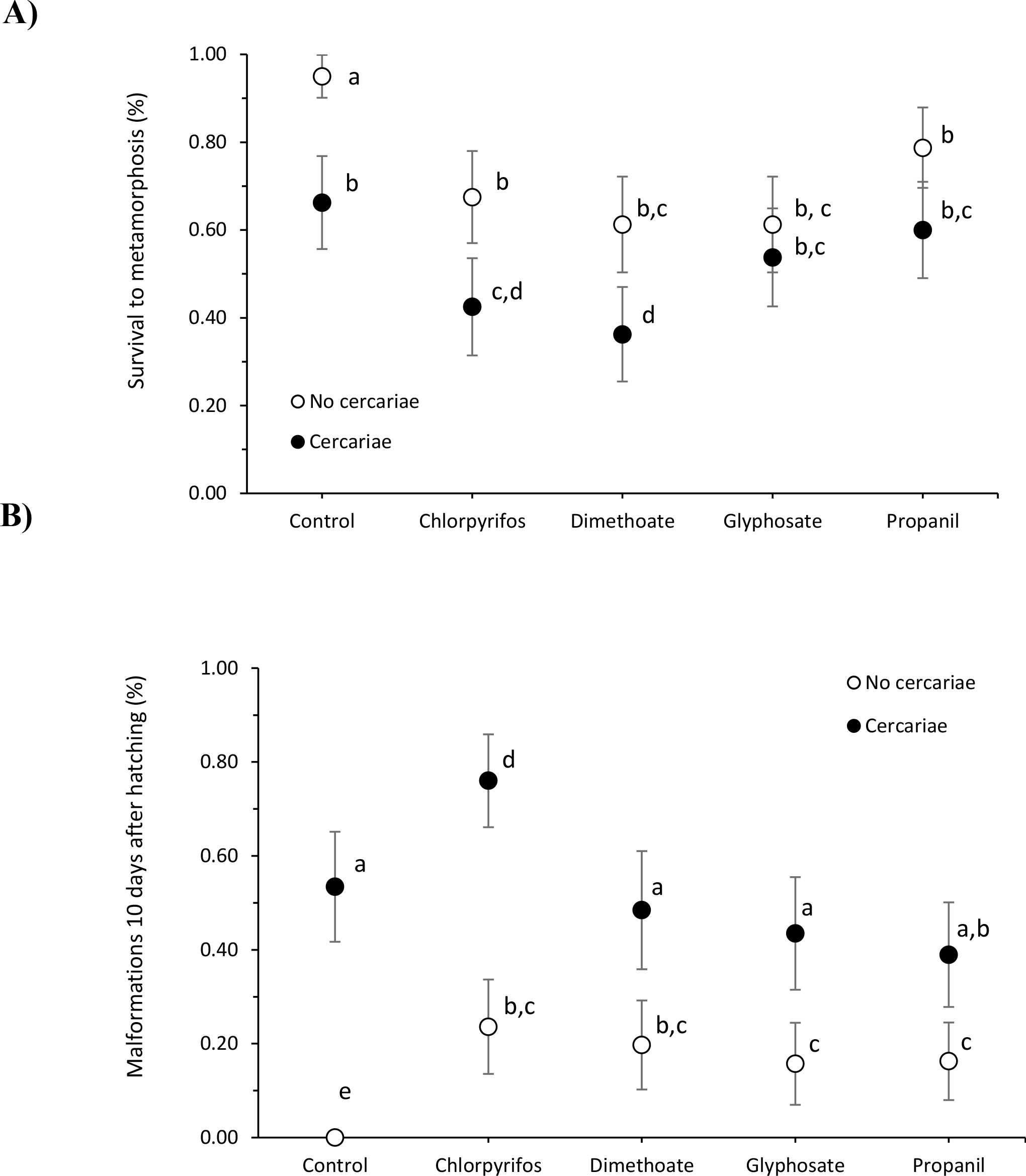
Mean proportion (± 95% confidence interval, *n* = 4 tanks) of *D. melanostictus* tadpoles that survived until metamorphosis (A) and that had malformations approximately 10 days post-hatching (B) after the exposure to five treatments (control dechlorinated water, chlorpyrifos, dimethoate, glyphosate and propanil) along with the presence or absence of exposure to cercariae of the trematode *Acanthostomum burminis*. Treatments that do not share letters were deemed significantly different from one another based on none overlapping confidence intervals.

### Malformations of tadpoles and metamorphs

Ten days after metamorphosis, tadpoles had significantly more malformations with than without pesticides (Main effect: χ^2^_4_= 40.11, *p*= 0.0001) and with than without cercariae (Main effect: χ^2^_1_= 138.33, *p*= 0.0001; Fig. 1B). None of the individuals in the absence of both cercariae and pesticides had malformations (Fig. 1B). In the absence of cercariae, chlorpyrifos, glyphosate, dimethoate, and propanil induced malformations in 24, 20, 16, and 16% of the tadpoles, respectively (Fig. 1B). In the absence of pesticides, cercariae induced malformations in 53% of the tadpoles (Fig.1B). Importantly, the effect of cercariae on malformation incidence depended on the pesticide treatment (Pesticide x cercariae: χ^2^_4_ = 28.10, *p*<0.001). Dimethoate, glyphosate, and propanil reduced the number of cercarial-induced malformations relative to the control and chlorpyrifos treatments (Fig. 1B). The malformations observed were scoliosis (vertebral column curvature, lateral deviation in the normally straight spine), kyphosis (hunched back, abnormal convexing of the spine), and edema.

### Size at metamorphosis

Despite affecting toad survival and malformations, we did not detect any effects of pesticide or cercarial treatments on body mass (Pesticide: *F*4,30=0.87, *p*=0.494, Cercariae: *F*1,30=0.48, *p* =0.492, Interaction: *F*4,30=0.24, *p* =0.914; Fig. 2A) or SVL at metamorphosis (Pesticide: *F*4,30=0.16, p =0.956, Cercariae: *F*1,30=0.42, *p* =0.520, Interaction: *F*4,30=1.32, *p* =0.284; Fig. 2B)

**Figure 2.**
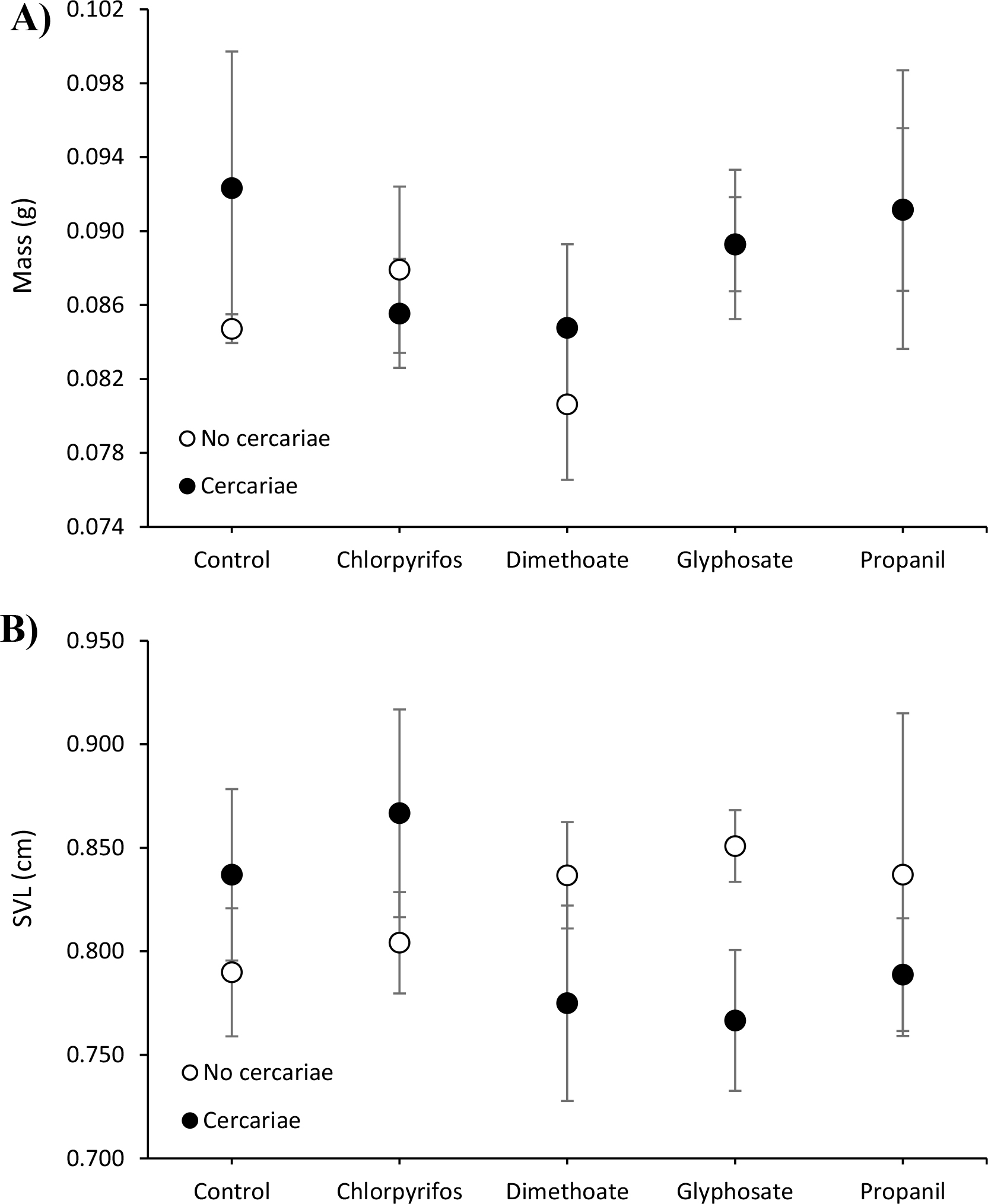
Mean (±SE, *n* = 4 tanks) mass (A) and snout-vent length (SVL) (B) of *D. melanostictus* toads at metamorphosis when exposed to five treatments (control dechlorinated water, chlorpyrifos, dimethoate, glyphosate and propanil) along with the presence or absence of exposure to cercariae of the trematode *Acanthostomum burminis*. There were no significant main effects of pesticides or cercariae on mass or SVL, nor was there a significant interaction between these predictors.

### Developmental rate

Days until 50% of the metamorphs had forelimb emergence and days to metamorphosis were significantly increased by exposure to pesticides (Main effect: *F*4,30=3.97, *p* =0.011; *F*4,30=4.28, *p* =0.007, respectively) and cercariae (Main effect: *F*1,30=12.48, *p* =0.001; *F*1,30=30.10, *p* <0.001, respectively, Fig. 3A, B). In the absence of cercariae, tadpoles exposed to chlorpyrifos, glyphosate, dimethoate, and propanil took 3.7, 5.1, 1.3, and 0.8 more days to metamorphose, respectively, relative to control tadpoles (Fig. 3B). In the absence of pesticides, tadpoles exposed to cercariae took 10.5 more days to metamorphose relative to those not exposed to cercariae (Fig. 3B). Additionally, the effect of cercariae on days until 50% of the frogs had forelimb emergence and days to metamorphosis depended on the pesticide treatment (Pesticide x cercariae: *F*4,30=2.95, *p*=0.036; *F*4,30=4.09, *p*=0.009, respectively). This was because cercariae increased and decreased development time in the presence of chlorpyrifos and dimethoate, respectively, relative to the pesticide control (Fig. 3A, B).

**Figure 3.**
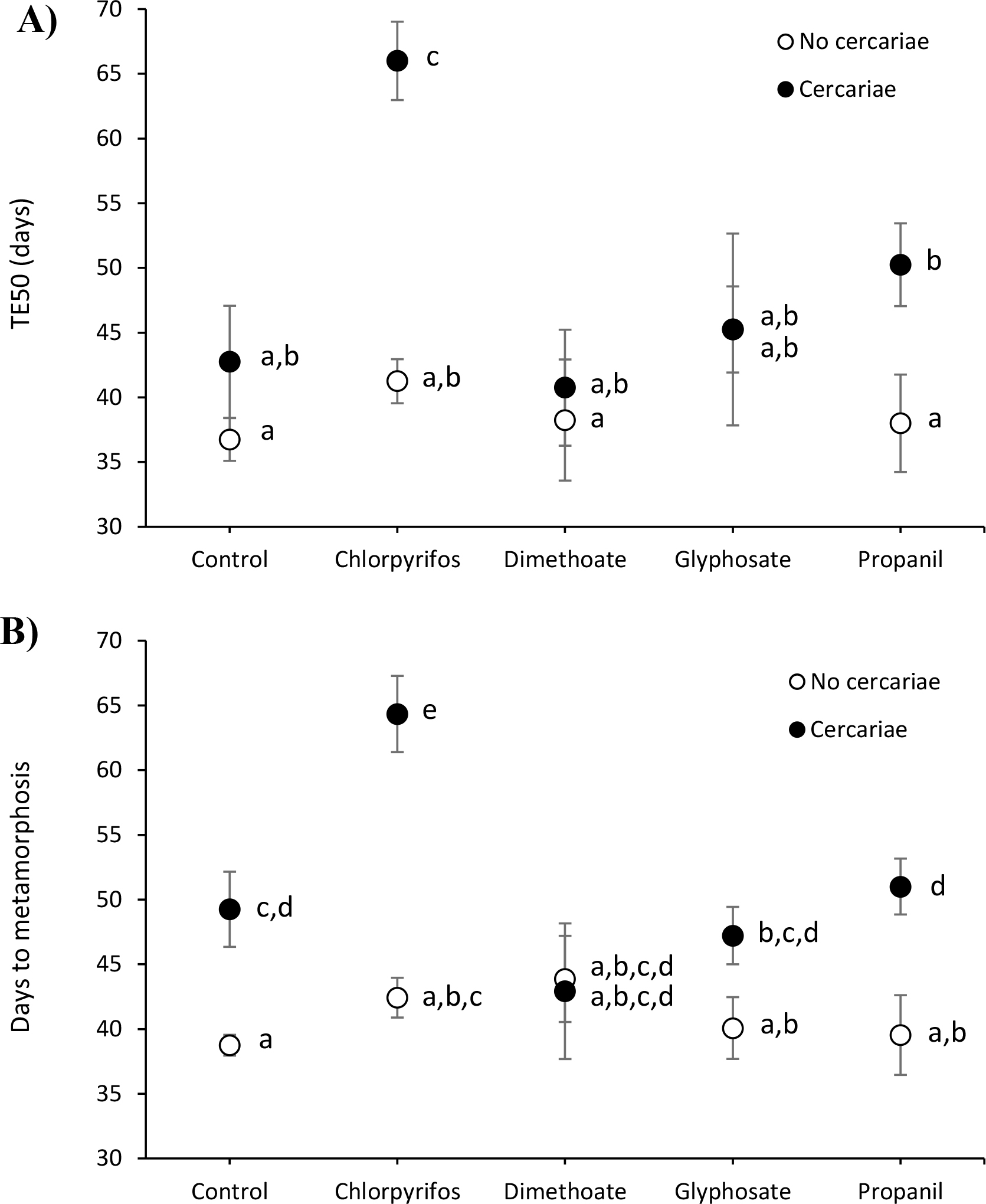
Mean experimental day when 50% of *D. melanostictus* toads had forelimb emergence (TE50) (A) and mean days to metamorphosis (**B**) when exposed to five treatments (control dechlorinated water, chlorpyrifos, dimethoate, glyphosate and propanil) crossed with the presence or absence of exposure to cercariae of the trematode *Acanthostomum burminis*. Treatments that do not share letters were deemed significantly different from one another based on a Fisher’s LSD post hoc multiple comparison test.

## DISCUSSION

Exposure to cercariae of *A. burminis* alone and the four pesticides alone significantly increased mortality and malformations in the Asian common toad compared to the water control. However, individual chemicals interacted with the parasites in different ways. Exposure to the cercariae in the presence of the two insecticides (chlorpyrifos and dimethoate) additively enhanced the effects on mortality induced by either treatment alone. However, exposure to the cercariae in the presence of the herbicides resulted in an antagonistic interaction where survival in the combined treatment was less than additive. Moreover, the effect of cercariae on malformation incidence depended on the pesticide treatment. Dimethoate, glyphosate, and propanil reduced the number of cercarial-induced malformations relative to the control and chlorpyrifos treatments.

In contrast to the current study, in a previous study on common hourglass tree frog tadpoles, the combined exposure of pesticides and cercariae resulted in a marked reduction in survival and significantly elevated levels of malformations compared to the lone exposures [45]. Differences in the traits of the Asian common toad and hourglass tree frog might explain these differences. The Asian common toad has thick, dry skin and the adults are nocturnal, terrestrial habitat generalists found frequently in human-altered agricultural and urban areas. The hourglass tree frog has thin skin and is an arboreal species found mostly in agricultural land, home gardens, houses, and other buildings. The differences in their skin are even visible at the tadpole stage, as tadpoles of the toad have thick dark skin and those of the frog have thin light skin.

Pesticide and cercarial treatments affected developmental traits of toads. There was no difference in the size of the tadpoles exposed to either cercariae or pesticides or both compared to the size of tadpoles in the water control. However, significant lengthening of the developmental period (i.e. days until 50% of the frogs had forelimb emergence and days to metamorphosis) was observed for tadpoles exposed to pesticides, cercariae, or both compared to the water control. Moreover, the effect of cercariae on the growth period depended on the pesticide treatment. Relative to the control, cercariae increased the developmental period in the presence of chlorpyrifos and decreased development in the presence dimethoate. In contrast to our results, Jayawardena et al. [45] discovered that the same combined cercarial and pesticide treatments as we used here significantly lengthened the growth period and reduced growth rates in the common hourglass tree frog relative to the two treatments alone.

The enhanced effect of pesticides on trematode disease severity might be due to impairment of the amphibian immune system. Immunosuppressive effects of pesticides have been reported in various studies [2, 40, 54–58]. Exact mechanisms of these immunosuppressive effects are unknown. However, Edge *et al*. [8] suggested that pesticides, particularly glyphosate, may affect skin peptides that can provide an important barrier to infections. Similarly, Gibble and Baer [59] reported that the in-vitro activity of antimicrobial peptides was reduced by agricultural runoff containing the herbicide atrazine [59]. In several other instances, pesticide exposure was associated with decreased melanomacrophage activity in the liver [60], reduced spleen cellularity [61], decreased lymphocyte proliferation [58], decreased white blood cell counts [62], and elevated parasite prevalence [41, 42, 56, 60]. On the other hand, pesticides may indirectly increase susceptibility to parasite infection by decreasing activity patterns [63] given that tadpoles can avoid free-swimming parasites by moving away from cercariae or swimming in erratic patterns [64, 65]. However, in the present study, tadpole ability to behaviorally avoid cercariae was controlled by exposing the tadpoles to parasites in a small volume of water where all the parasites successfully penetrated the tadpole, forcing them to rely primarily on physiological defenses, such as immune responses [66].

None of the pesticides tested in the present study had any detrimental effect on trematode survival. Raffel *et al*. [66] measured the effects of atrazine, glyphosate, carbaryl, and malathion on embryo and miracidium (free-living stage) survival of *Echinostoma trivolvis*, a common trematode of amphibians, and found no evidence of effects of these pesticides at ecologically relevant concentrations. In addition, the survival of renicolid cercariae improved with increasing concentrations of the common herbicide glyphosate, with cercariae living about 50% longer in mg a.i. L^-1^ of glyphosate than in control conditions [66]. In addition, several studies [e.g., 70-74] have examined the effect of pollutants on either the output of cercariae from snails or their subsequent survival. For instance, Kelly *et al*. [60] recently showed that the New Zealand snail *Potamopyrgus antipodarum* released approximately three times more *Telogaster opisthorchis* cercariae per day when exposed to glyphosate than when kept in glyphosate-free water. In many cases, exposure to pollutants, such as metals, pesticides, and herbicides, reduces replication of trematodes within snails [67, 68] or their rate of emergence from snails [46, 69]. However, some studies report reduced virulence of trematode infections in the presence of chemicals. For instance, Koprivnikar *et al*. [70] showed that trematode cercariae exposed to atrazine has less success in infesting the tadpoles than those in the control groups.

As described by Rohr *et al*. [4], the majority of *Acanthostomum* cercariae crawl towards the cloacal vent and form cysts in the crease between the body and tail, where limb buds are located. However, *A. burminis* does not appear to be as virulent as the more well-known *Ribeiroia ondatrae* that also causes amphibian limb deformities. Unlike *Ribeiroia*, *Acanthostomum* cysts are not visible as swollen lumps, perhaps because of their smaller size [71]. Apparently, *Ribeiroia* cysts average 300-350 μm in length (excysted metacercariae, 500-650 μm and adult 4160-5250 μm in length; [71]), whereas *Acanthostomum* cercariae average 216 μm in length. Size of cercariae has been suggested to affect the virulence of trematode infections [4], with larger metacercariae presumably causing more tissue damage, eliciting greater immune responses, and consuming more host resources.

In the field, combinations of pesticides and trematodes may have adverse or beneficial effects on amphibian populations. Pesticides may enhance snail population densities or immunosuppress hosts, thereby promoting deadly amphibian infections [32, 41, 60]. Mesocosm studies conducted by Rohr and colleagues [34] revealed that atrazine increases algal and snail biomass and increases trematode loads in immunosuppressed *Rana pipiens* tadpoles. In the present study, exposures to cercariae in the presence of the two insecticides further reduced host survival relative to the cercariae or insecticides alone. In contrast, herbicides had less than additive effects on mortality associated with cercarial exposures.

In many instances, pesticide concentrations in waterbodies are too low to cause direct amphibian mortality. However, their interactions with other biotic and abiotic factors can induce substantial amphibian mortality, as shown in the current study. Hence, the effects of multiple stressors must be more thoroughly considered in ecological risk assessments of wildlife [72].

## List of abbreviations

General linear model (GLM), Snout vent length (SVL), Time to metamorphosis (TE50)

## Declarations

### Ethics approval and consent to participate

Authors declare that the experiments conducted, complied with the current laws of Sri Lanka. Approval was obtained for the collection of wildlife specimens from the protected areas, and for conducting animal research, from the Department of Wildlife Conservation, Sri Lanka and from the Ethics Review Committee, Postgraduate Institute of Science, University of Peradeniya, respectively. Hence, all experiments conducted were in compliance with ethical guidelines provided by these two authorities.

### Consent for publication

Not applicable

### Availability of data and material

The data sets supporting the results of this article are included within the article and its additional files.

### Authors’ contributions

UA carried out the study under the guidance of RS, AN, and PH. JRR guided in analyses and interpreting the results. UA drafted the manuscript and RS, JRR and PH reviewed it before the initial submission. All authors read and approved the final manuscript.

## Acknowledgements

Authors thank V. Imbuldeniya and Y.G. Ariyaratne of the Department of Zoology, University of Peradeniya for technical assistance.

